# Backbone Assignment of a 28.5 kDa Class A Extended Spectrum β-Lactamase by High-Field, Carbon-Detected Solid-State NMR

**DOI:** 10.1101/2025.05.18.654753

**Authors:** Christopher G. Williams, Songlin Wang, Alexander F. Thome, Owen A. Warmuth, Varun Sakhrani, Chad M. Rienstra, Leonard J. Mueller

## Abstract

^13^C and ^15^N backbone chemical shift assignments are reported for the 28.5 kDa protein Toho-1 β-lactamase, a Class A extended spectrum β-lactamase. A very high level of assignment completeness (97% of the backbone) is enabled by the combined sensitivity and resolution gains of ultrahigh-field NMR spectroscopy (1.1 GHz), improved probe technology, and optimized pulse sequences. The assigned chemical shifts agree well with our previous solution-state NMR assignments, indicating that the secondary structure is conserved in the solid state. These assignments provide a foundation for future investigations of sidechain chemical shifts and catalytic mechanism.

## 1. Biological Context

Since the introduction of penicillin, the widespread use of β-lactam antibiotics has driven the emergence of antibiotic resistance through the evolution of β-lactamases – enzymes that inactivate these drugs by hydrolyzing their therapeutically active β-lactam ring.^1,2^ These enzymes contribute to resistance even against some of the most effective antibiotics. Of the four classes of β-lactamases, Classes A, C, and D utilize a catalytic serine in the active site, while the Class B metallo-beta-lactamases rely on a zinc ion for hydrolysis. Despite extensive studies aimed at elucidating enzyme mechanism and function, key questions remain, particularly regarding the roles of the active site residues.^3^ Within Class A β-lactamases, studies have identified mechanistic involvement of residues Ser70, Lys73, Glu166, and an active site water molecule, while also implicating nearby residues such as Ser130 and Lys234, although their precise roles remain unclear.^4-8^

A major challenge in combating antibiotic resistance is posed by extended-spectrum β-lactamases (ESBL), which exhibit increased activity towards first, second, and third generation cephalosporins, as well as monobactam antibiotics.^2,9,10^ One such Class A ESBL is Toho-1 β-Lactamase (Toho-1; also known as CTX-M-44), a 263-residue enzyme with greater activity towards cefotaxime than other oxyimino-beta-lactam substrates such as cefepime, ceftriaxone, and ceftazidime. The CTX-M class is now the most prevalent group of ESBLs worldwide.^11^ Gaining a deeper mechanistic understanding of these enzymes, and their interactions with both antibiotics and inhibitors, is therefore vital for the development of effective antimicrobial therapies.

Our initial studies of Toho-1 using solution-state NMR provided backbone assignments for both its free and inhibitor-bound forms, along with preliminary dynamics data that laid the foundation for understanding how the backbone dynamics relate to inhibitor binding.^12^ A critical next step is to use NMR spectroscopy, with its site-specific resolution, to probe the chemistry occurring in the active site – an effort that requires detecting signals from amino acid sidechains. However, β-lactamases, with molecular weights typically around 30 kDa, approach the practical size limit for detecting complete sets of sidechain resonances using solution-state NMR, making it particularly challenging to access detailed information about the catalytic residues.^13,14^

This work serves as an important step toward the goal of expanding the chemical assignments for Toho-1. Our approach relies on recent advances in solid-state NMR (SSNMR), which provides an alternate route to accessing sidechains. In SSNMR, the protein is immobilized within the crystal lattice and dipolar line narrowing is achieved by magic-angle spinning (MAS), which does not rely on molecular tumbling.^15,16^ At high magnetic field, spectral linewidths for ^13^C-detected MAS spectra rival the resolution observed in solution, and the coherence lifetimes are sufficiently long to enable multiple magnetization transfers and the acquisition of highly resolved 3D spectra to assign increasingly larger proteins such as Toho-1.^17-20^ Our approach combines ultrahigh field NMR at 1.1 GHz with long-observation-window band-selective homonuclear decoupling (LOW-BASHD), alongside recent improvements in probe, receiver, external ^2^H lock, and ^1^H decoupling performance, to achieve significantly enhanced sensitivity and resolution in carbon-detected biomolecular SSNMR.^21,22^ The backbone assignments serve as an essential first step toward accessing complete sidechain resonances. These assignments also enable an evaluation of the extent to which SSNMR results for Toho-1 align with those obtained in solution, thereby providing a bridge between prior solution-state studies and future solid-state investigations of β-lactamases.

## 2. Methods and experiments

### 2.1 Protein Expression, Purification, and Crystallization

Uniformly-^13^C,^15^N labeled Toho-1 was expressed and purified following our previously published procedure with slight modifications.^12,23,24^ In brief, BL21 *E. coli* cells infused with a Toho-1 expression plasmid were cultured overnight in kanamycin-supplemented LB broth in a shaking incubator set to 250 rpm and 303 K. Subsequently, 20 ml of this culture was added to

500 ml of kanamycin-supplemented LB broth, equally divided between two 2.8 L baffled Fernbach flasks and incubated an additional 6 hrs until an OD_600_ of 1.9 was reached. At this stage, the cells were pelleted using a room temperature floor centrifuge and resuspended in 500 mL of kanamycin-supplemented minimal media containing 4.0 g ^13^C-glucose and 3.0 g ^15^N-NH_4_Cl, evenly split between two 2.8 L baffled Fernbach flasks. The flasks were placed in a shaking incubator at 250 rpm and room temperature for 15 minutes before sorbitol and betaine were added at a final concentration of 200 mM and 5 mM, respectively, and allowed to incubate for another 15 minutes. Finally, IPTG was added to a final concentration of 1 mM and the cells grown overnight. The following morning, cell harvesting was performed through centrifugation, and the cells were resuspended in 20 mM MES buffer at pH 6.5 (Buffer A). The iced mixture was then lysed using a Branson 450D Digital Sonifier (Emerson Industrial Automation, St. Louis, MO, USA), and the cell debris was removed by centrifugation. The resulting supernatant was collected, filtered through Kimwipes, diluted with Buffer A, and loaded onto a 5mL HiTrap SP Sepharose FF column (GE Healthcare, Pittsburg, PA, USA) equilibrated with Buffer A. Protein elution was carried out utilizing a linear gradient of 20mM MES buffer at pH 6.5 and containing 300mM NaCl and tracked via UV absorbance. Eluted protein fractions were pooled and further purified using a 120mL HiLoad Superdex S-200 SEC column equilibrated with Buffer A. Protein elution was tracked by UV absorbance, and fractions containing the target protein were pooled and concentrated using a 10 kDa MWCO Ultra Centrifugal Filter (Amicon) to a concentration of 300-400 μM.

Protein microcrystals were grown by mixing the protein at 300-400 μM in Buffer A with a crystallization buffer containing 7 mM spermine and 30% PEG-8k in Buffer A at a 1:1 ratio. Crystals were allowed to grow over 3-5 days at 4° C. The sample was packed into a 1.6 mm Varian style SSNMR rotor using customized sample packing devices.^25,26^

### 2.2 Solid-State NMR Spectroscopy

Solid-state NMR spectra were collected at NMRFAM on a Bruker NEO (1.1 GHz) spectrometer using a Black Fox (Tallahassee, FL) triple resonance probe in HCN mode. The probe incorporates a PhoenixNMR (Loveland, CO) 1.6 mm spinning module and has a dual coil design, with an inner solenoid tuned to ^13^C/^15^N and an outer low-electric-field ^1^H resonator.^27,28^ All experiments were conducted under 25 kHz MAS and were optimized using a combination of manual and automated methods, employing strategies for stable CP as described.^29,30^

#### 2.2.1 ^15^N/^13^C_α_ 2D correlation experiment (NCA)

The 2D NCA SPECIFIC CP experiment was performed with the ^13^C carrier frequency set at 55 ppm.^31^ The ^15^N polarization was prepared via an adiabatic CP with a downward tangential ramp pulse applied on the ^1^H channel, with a contact time of 2.0 ms, ^15^N RF amplitude of 40 kHz, and an average ^1^H RF amplitude of 60 kHz.^32^ ^15^N polarization was then transferred to ^13^C_α_ using CP with an upward tangential ramp on the ^13^C channel, with a contact time of 7.0 ms, ^15^N RF amplitude of 10 kHz, and an average ^13^C RF amplitude of 15 kHz. CW decoupling of ^1^H at 100 kHz was applied during this period. The t_1_ acquisition time was 32 ms with a 160 µs increment for 200 complex points, while a 5.2 µs ^13^C π-pulse was applied at the center of the t_1_ period to decouple ^*1*^*J*_*NC*_. The t_2_ acquisition time was 30.7 ms with a 10 µs dwell time for 3072 complex time-domain points. SPINAL-64 ^1^H decoupling at 100 kHz was applied during acquisition.^33^ LOW-BASHD detection was implemented in the direct dimension to decouple the ^*1*^*J*_*CαC′*_ using τ_dec_=5.0 ms and 72.5 µs cosine modulated Gaussian π-pulses, with RF amplitude at 19 kHz and frequency set at 175 ppm.^21^ The recycle delay was 1.5 s. The total experimental time was 5.3 hours.

#### 2.2.2 ^15^N/^13^CO 2D correlation experiment (NCO)

The 2D NCO SPECIFIC CP experiment was performed with the ^13^C carrier frequency set at 175 ppm.^31^ The ^15^N polarization was prepared via an adiabatic CP with a downward tangential ramp pulse applied on the ^1^H channel, with a contact time of 2.0 ms, ^15^N RF amplitude of 40 kHz, and an average ^1^H RF amplitude of 60 kHz.^32^ ^15^N polarization was then transferred to ^13^CO using CP with an upward tangential ramp on the ^13^C channel, with a contact time of 7.0 ms, ^15^N RF amplitude of 16 kHz, and an average ^13^C RF amplitude of 10 kHz. CW decoupling of ^1^H at 100 kHz was applied during this period. The t_1_ acquisition time was 32 ms with a 160 µs increment for 200 complex points, while a 5.2 µs ^13^C π-pulse was applied at the center of the t_1_ period to decouple ^*1*^*J*_*NC*_. The t_2_ acquisition time was 30.7 ms with a 10 µs dwell time for 3072 complex points. SPINAL-64 ^1^H decoupling at 100 kHz was applied during acquisition.^33^ LOW-BASHD detection was implemented in the direct dimension to decouple the ^*1*^*J*_*CαC′*_ using τ _dec_ =5.0 ms and72.5 µs cosine modulated Gaussian π-pulses, with RF amplitude at 19 kHz and frequency set at 55 ppm.^21^ The recycle delay was 1.5 s. The total experimental time was 5.3 hours

#### 2.2.3 ^15^N/^13^C_α_/^13^CO 3D correlation experiment (NCACO)

The 3D NCACO experiment was performed with the ^13^C carrier frequency set at 55 ppm. The ^1^H to ^15^N CP and the ^15^N to ^13^C_α_ CP transfer conditions were identical to the 2D NCA experiment as described above. ^13^C_α_ to ^13^CO polarization transfer was achieved using a RF-driven Dipolar Recoupling (RFDR) mixing.^34^ The contact time was 1.28 ms, with a ^13^C RF amplitude of 135 kHz. CW ^1^H decoupling at 100 kHz was applied during this RFDR mixing period. The t_1_ acquisition time was 20.0 ms with a 200 µs increment for 100 complex points, and a 5.2 µs ^13^C π-pulse was applied at the center of the t_1_ period to decouple ^*1*^*J*_*NC*_. The t_2_ acquisition time was 7.2 ms with an 80 µs increment for 90 complex points. At the center of the t_2_ period, a 5.2 µs ^13^C hard π-pulse, a 300 µs ^13^C soft π-pulse with RSNOB shape at 55 ppm, and a 14.8 µs ^15^N π-pulse were applied to decouple the ^*1*^*J*_*CαCX*_ and ^*1*^*J*_*NC*_.^35^ The t_3_ acquisition time was 30.7 ms with a 10 µs dwell time for 3072 complex points. SPINAL-64 ^1^H decoupling at 100 kHz was applied during acquisition.^33^ LOW-BASHD detection was implemented in the direct dimension to decouple the ^*1*^*J*_*CαC′*_ using τ_dec_ =5.0 ms and 72.5 µs cosine modulated Gaussian π-pulses, with RF amplitude at 19 kHz and frequency set at 55 ppm.^21^ The recycle delay was 1.5 s. The 3D spectrum was acquired using a 25% NUS schedule, and the total experimental time was 15.4 hours.

#### 2.2.4 ^15^N/^13^CO/^13^C_α_ 3D correlation experiment (NCOCA)

The 3D NCOCA experiment was performed with the ^13^C carrier frequency set at 175 ppm. The ^1^H to ^15^N CP and the ^15^N to ^13^CO CP transfer conditions were identical to the 2D NCO experiment as described above. The ^13^CO to ^13^C_α_ polarization transfer was achieved using 1.28 ms RFDR mixing with a ^13^C RF amplitude of 135 kHz.^34^ CW ^1^H decoupling at 100 kHz was applied during this RFDR mixing period. The t_1_ acquisition time was 20.0 ms with a 200 µs increment for 100 complex points. A 5.2 µs ^13^C π-pulse was applied at the center of the t_1_ period to decouple the ^*1*^*J*_*NC*_. The t_2_ acquisition time was 7.7 ms with a 160 µs increment for 48 complex points. At the center of the t_2_ period, a 5.2 µs ^13^C hard π-pulse, a 300 µs ^13^C soft π-pulse with RSNOB shape at 175 ppm, and a 14.8 µs ^15^N π-pulse were applied to decouple the ^*1*^*J*_*CαCX*_ and ^*1*^*J*_*NC*_.^35^ The t_3_ acquisition time was 30.7 ms with a 10 µs dwell time for 3072 complex time-domain points. SPINAL-64 ^1^H decoupling at 100 kHz was applied during acquisition.^33^ LOW-BASHD detection was implemented in the direct dimension to decouple the ^*1*^*J*_*CαC’*_ using τ_dec_ =5.0 ms and 72.5 µs cosine modulated Gaussian π-pulses, with RF amplitude at 19 kHz and frequency set at 175 ppm.^21^ The recycle delay was 1.5 s. The 3D spectrum was acquired using a 25% NUS schedule and the total experimental time was 33.5 hours.

#### 2.2.5 ^13^C_α_/^15^N/^13^CO 3D correlation experiment (CANCO)

The 3D CANCO experiment was performed with the ^13^C carrier frequency set to 175 ppm. ^13^C_α_ polarization was prepared via adiabatic CP using a downward tangential ramp pulse on the ^1^H channel, with the ^13^C frequency adjusted to 55 ppm for ^1^H-^13^C_α_ transfer.^32^ The CP contact time was 1.5 ms, with a ^13^C RF amplitude of 108 kHz, and an average ^1^H RF amplitude of 81 kHz. Polarization was then transferred from ^13^C_α_ to ^15^N via CP using an upward tangential ramp on the ^13^C channel. The contact time was 7.0 ms, ^15^N RF amplitude of 10 kHz, and an average ^13^C RF amplitude was 15 kHz. CW ^1^H decoupling at 100 kHz was applied during this CP period. The ^15^N polarization was transferred to ^13^CO using CP with an upward tangential ramp on the ^13^C channel after setting the ^13^C frequency back to 175 ppm. The contact time was 7.0 ms, with a ^15^N RF amplitude of 10 kHz, and an average ^13^C RF amplitude of 16 kHz. CW ^1^H decoupling at 100 kHz was applied during this CP period. The t_1_ acquisition time was 7.2 ms with an 80 µs increment for 90 complex points. A 5.2 µs ^13^C hard π-pulse, a 300 µs ^13^C soft π-pulse with RSNOB shape at 55 ppm, and a 14.8 µs ^15^N π-pulse were applied at the center of the t_1_ period to decouple the ^*1*^*J*_*CαCX*_ and ^*1*^*J*_*NC*_.^35^ The t_2_ acquisition time was 20.0 ms, with a 200 µs increment for 100 complex points. A 5.2 µs ^13^C π-pulse was applied at the center of the t_2_ period to decouple the ^*1*^*J*_*NC*_. The t_3_ acquisition time was 30.7 ms, with a 10 µs dwell time for 3072 complex time-domain points. SPINAL-64 ^1^H decoupling at 100 kHz was used during acquisition.^33^ LOW-BASHD detection was implemented in the direct dimension to decouple the ^*1*^*J*_*CαC′*_ using τ_dec_ =5.0 ms and 72.5 µs cosine modulated Gaussian π-pulses, with RF amplitude at 19 kHz and frequency set at 55 ppm.^21^ The recycle delay was 1.5 s. The 3D spectrum was acquired using a 25% NUS schedule, and the total experimental time was 30.8 hours.

#### 2.2.6 ^13^C_β_/^13^C_α_/^13^CO 3D correlation experiment (CBCACO)

The 3D CBCACO experiment was performed with the ^13^C carrier frequency set at 40 ppm. ^13^C_β_ polarization was prepared via adiabatic CP using a downward tangential ramp pulse on the ^1^H channel. The CP contact time was 2.5 ms, with a ^13^C RF amplitude of 108 kHz, and an average ^1^H RF amplitude of 82 kHz. Polarization was then transferred from ^13^C_β_ to ^13^C_α_ via Dipolar Recoupling Enhancement through Amplitude Modulation (DREAM) using a downward tangential ramp on the ^13^C channel.^36^ The contact time was 3.0 ms, and an average ^13^C RF amplitude was 12 kHz. CW ^1^H decoupling at 100 kHz was applied during this DREAM mixing period. The ^13^C_α_ polarization was transferred to ^13^CO using RFDR mixing.^34^ The contact time was 1.28 ms, with a ^13^C RF amplitude of 135 kHz. CW ^1^H decoupling at 100 kHz was applied during this RFDR mixing period. The t_1_ acquisition time was 6.4 ms with a 40 µs increment for 160 complex points. The t_2_ acquisition time was 6.4 ms with an 80 µs increment for 80 complex points. At the center of the t_2_ period, a 5.2 µs ^13^C hard π-pulse, a 300 µs ^13^C soft π-pulse with RSNOB shape at 55 ppm, and a 14.8 µs ^15^N π-pulse were applied to decouple the ^*1*^*J*_*CαCX*_ and ^*1*^*J*_*NC*_ .^35^ The t_3_ acquisition time was 30.7 ms with a 10 µs dwell time for 3072 complex time-domain points. SPINAL-64 ^1^H decoupling at 100 kHz was applied during acquisition.^33^ LOW-BASHD detection was implemented in the direct dimension to decouple the ^*1*^*J*_*CαC′*_ using τ_dec_ =5.0 ms and 72.5 µs cosine modulated Gaussian π-pulses, with RF amplitude at 19 kHz and frequency set at 55 ppm.^21^ The recycle delay was 1.5 s. The 3D spectrum was acquired using a 25% NUS schedule, and the total experimental time was 43.8 hours.

## 3. Results and Discussion

### 3.1 Solid-State NMR Backbone Assignments and Data Deposition

Protein backbone assignment of U-^13^C,^15^N-Toho-1 were conducted in CCPNmr AnalysisAssign.^37^ The backbone ^13^C and ^15^N chemical shifts have been deposited at the Biological Magnetic Resonance Bank (BMRB) database (http://www.bmrb.wisc.edu) under accession number 53038.^38^

### 3.2 Assignments and Completeness

Data collection followed optimized SSNMR protocols for backbone assignments and consisted of experiments to establish both the intra-residue (2D NCA; 3D NCACX and CBCACO) and inter-residue (2D NCO; 3D NCOCX and CANCO) correlations essential for the backbone walk.^39^

Figure 1 highlights the exceptional resolution achieved for this microcrystalline protein in the 2D NCA correlation spectrum acquired at 1.1 GHz and a MAS rate of 25 kHz. Effective homonuclear decoupling in both the indirect and direct dimensions is essential to obtain such high resolution. To accomplish this, LOW-BASHD was employed to remove the ∼55 Hz *J*_CαC’_ splitting in the directly detected ^13^C dimension.^21^ Indeed, the high degree of crystalline order in this sample enables resolution in which the fine structure due to homonuclear *J*-couplings can routinely be observed and, in many cases, becomes the primary source of inhomogeneous line broadening.

**Figure 1:**
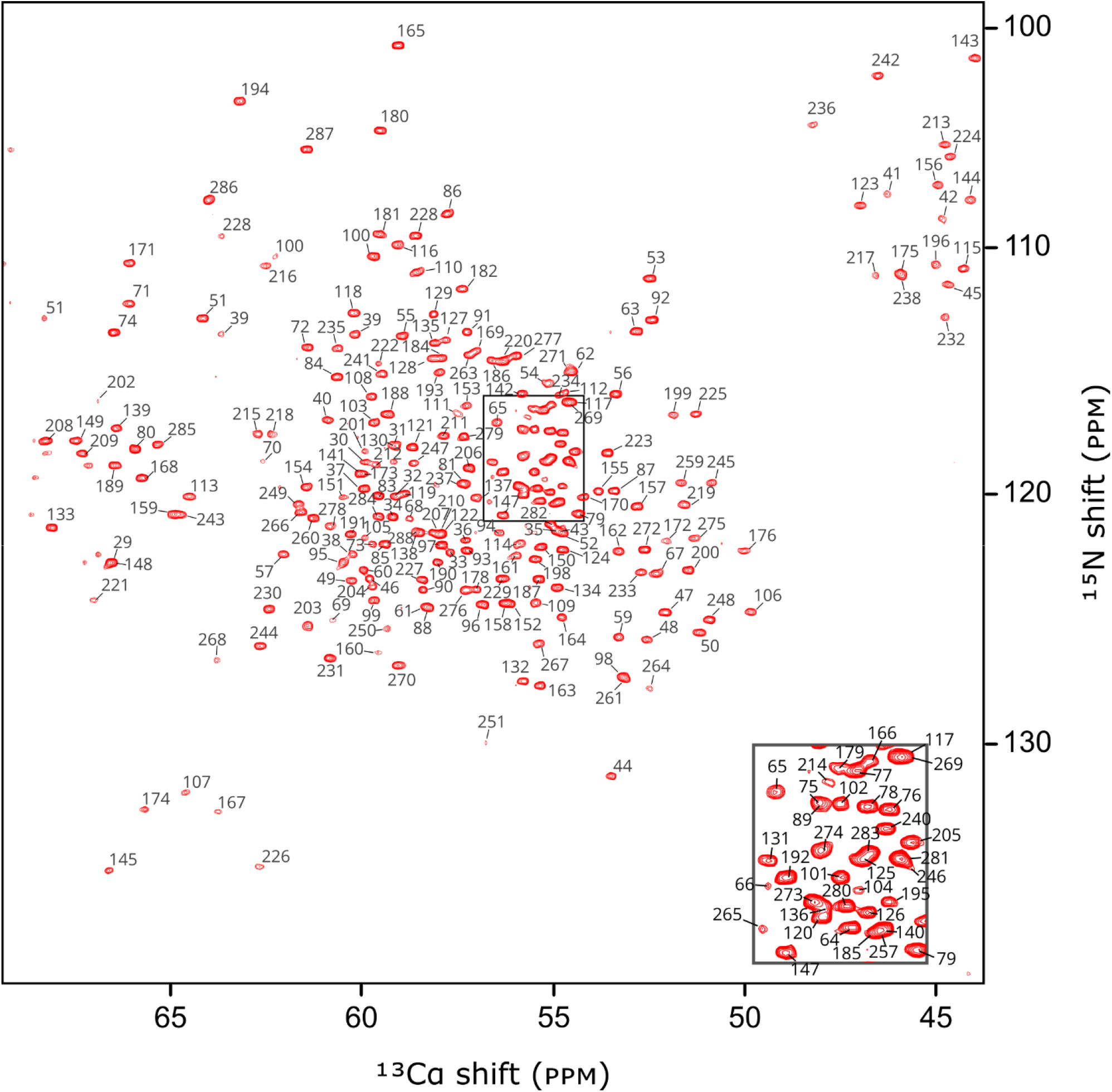
The 2D NCA correlation spectrum of U-^13^C,^15^N-labeled Toho-1 β-lactamase acquired at 1.1GHz (^1^H) under 25 kHz MAS. The spectrum demonstrates the high resolution achievable for this microcrystalline protein sample. LOW-BASHD decoupling was used to remove the ∼55 Hz ^1^*J*_*CαC′*_ splitting in the directly detected ^13^C dimension.^19^

The combined use of NCACO, CANCO, and NCOCA spectra enables a clear and continuous backbone walk linking intra- and inter-residue connectivity, as illustrated in Figure 2. While some C_β_ resonances can be found in the NCOCA spectrum, more comprehensive coverage was achieved through the CBCACO experiment, resulting in 227 out of 243 (93%) assigned C_β_ shifts. As in our previous solution-state study, the primary limitations in completeness stem from a more dynamic region of the protein spanning residues 252-257.^12^ However, overall backbone assignment completeness is high, with 992 of 1026 (97%) shifts assigned, as summarized in Figure 3.

**Figure 2:**
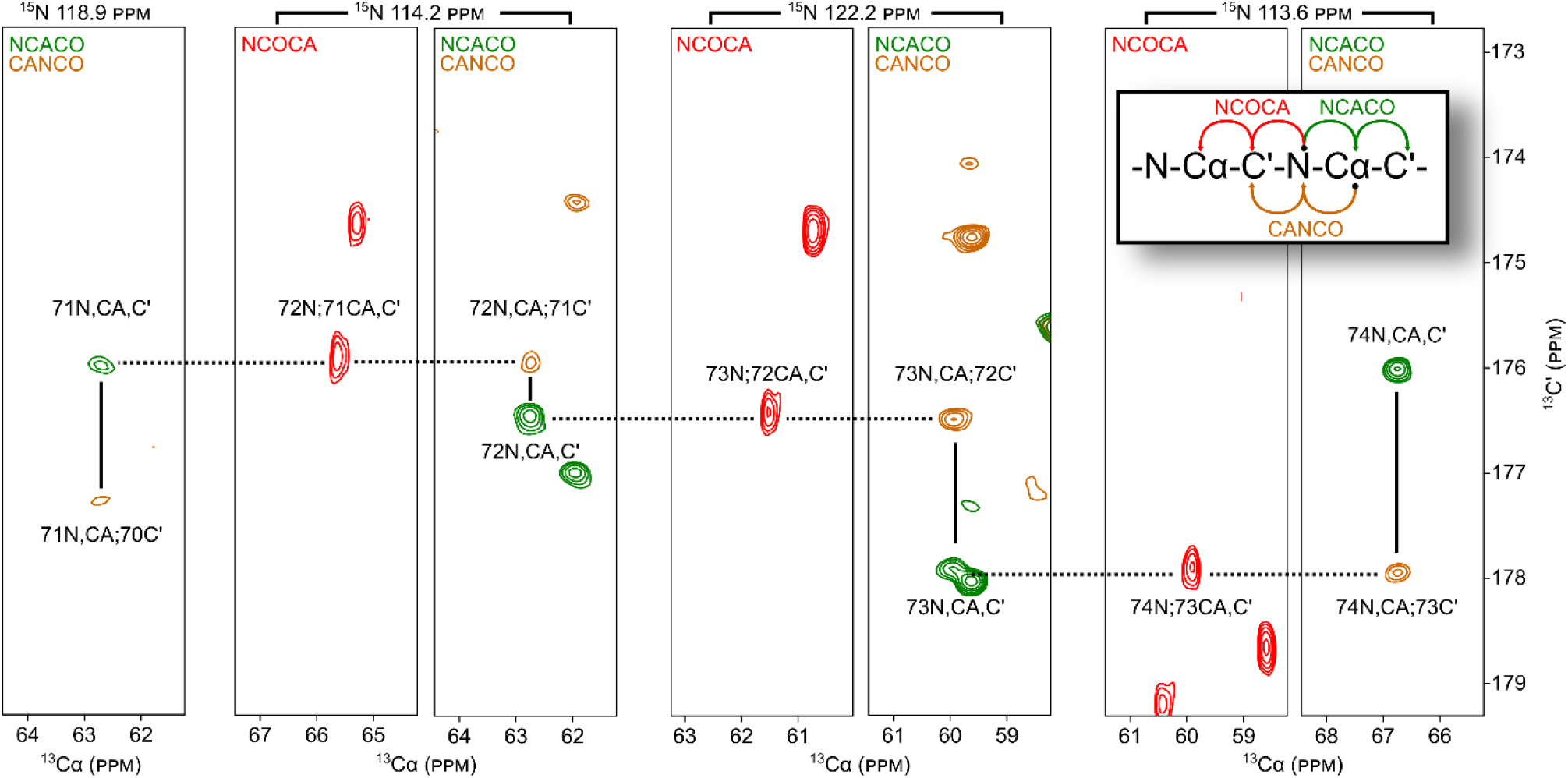
Two-dimensional strip plots illustrating a backbone walk from Val74 to Ser70 (including catalytic residues Ser70 and Lys73) using 3D NCACO (green), 3D CANCO (gold), and 3D NCOCA (red) spectra. Solid lines indicate correlations observed within the same NCA plane, corresponding to the carbonyl carbon resonances of sequential residues (i and i−1). Dashed lines trace the connection of the carbonyl carbon of residue i−1, detected via the NCOCA experiment, back to its own NCA plane in the NCACO spectrum. The amide nitrogen chemical shift for each plane is labeled above the corresponding strip.

**Figure 3:**
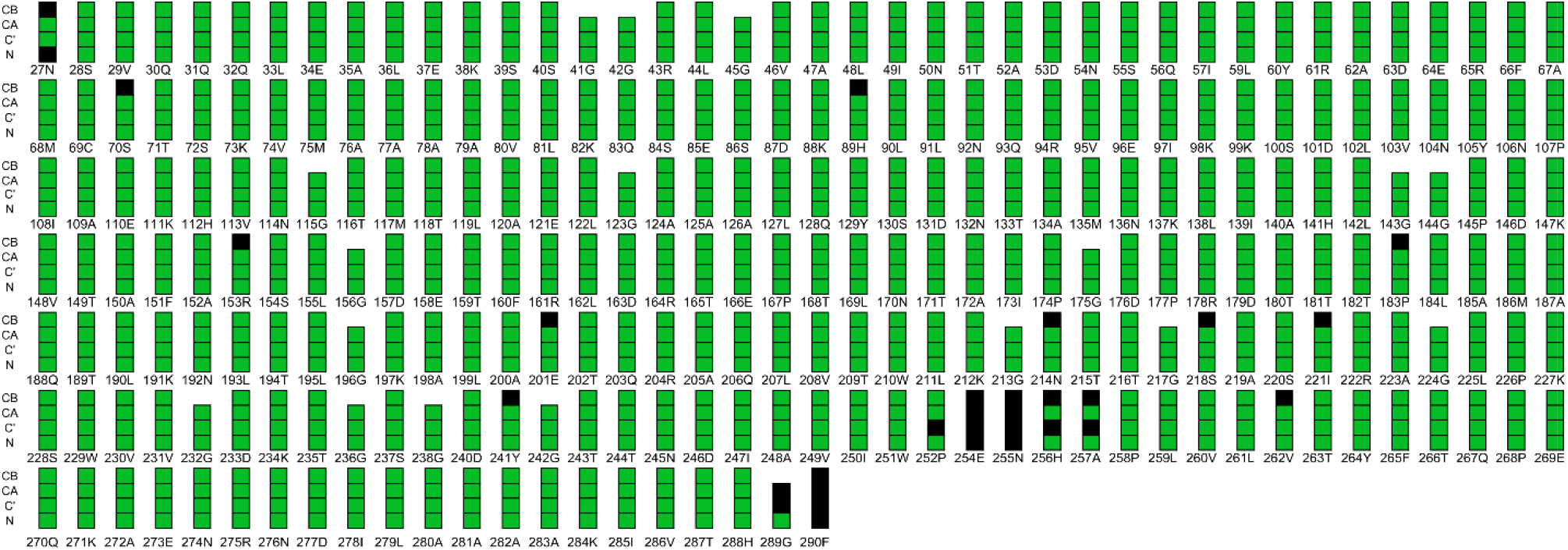
Backbone assignment completeness for Toho-1. Green and black boxes represent assigned and unassigned resonances respectively.

### 3.3 Active Site Chemical Shifts

Our previous solution-state NMR study revealed notable behavior regarding Ser70: both its N-H correlation and that for the substrate coordinating residue Ser237 were absent in the ^15^N HSQC spectrum of the free enzyme but appeared upon ligand binding.^12^ This behavior was attributed to dynamic signal broadening that stabilized upon substrate binding.^24,40^ Interestingly, in the current SSNMR experiments, backbone signals for Ser70 are detected, although not for its sidechain C_β_ resonance. We attribute this to an unfavorable dynamic regime, as supported by the absence of any Ser70 peak in the CBCACO spectrum, as shown in Figure 4. Encouragingly, backbone chemical shifts for the catalytically important Lys73 and Glu166 are observed, providing access to the active site and laying the groundwork for future sidechain-specific analysis.

**Figure 4:**
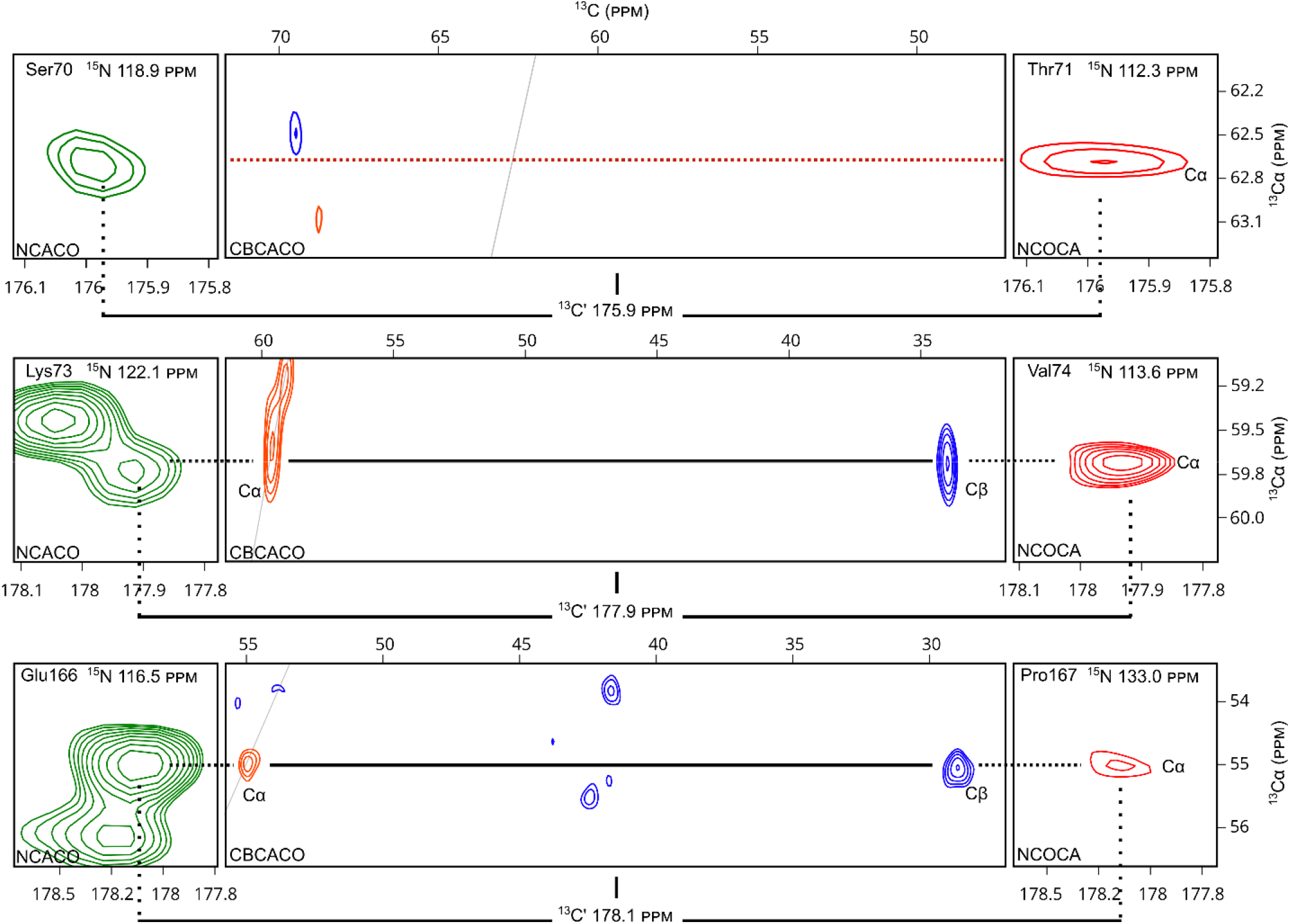
Backbone chemical shift correlations for active-site residues Ser70, Lys73, and Glu166 shown within NCACO (green), CBCACO (orange and blue), and NCOCA (red) 3D correlation spectra. Intra-residue correlations are observed within NCACO and CBCACO spectra, while inter-residue correlations are identified in the NCOCA strips from the nitrogen of their *i*+1 neighbor. Solid black lines match each residue’s C_α_–C′ resonance across the three spectra. The dashed red line highlights the absence of C_α_ and C_β_ peaks for Ser70 in the CBCACO spectrum.

### 3.4 Comparison to Solution State

Comparison with our previously published solution-state NMR assignments (Figure 5) reveals a close correspondence between datasets and supports the structural consistency of Toho-1 in the solution and crystalline states. Several outliers noted correspond to residues located in or near regions of increased conformational dynamics in solution.^12^ The correspondence between the solution and solid-state chemical shifts is essential for validating future solid-state analyses of Toho-1 and ensures that conclusions drawn from solid-state experiments can be meaningfully compared to, and integrated with, studies of related enzymes characterized in solution.^41-46^

**Figure 5:**
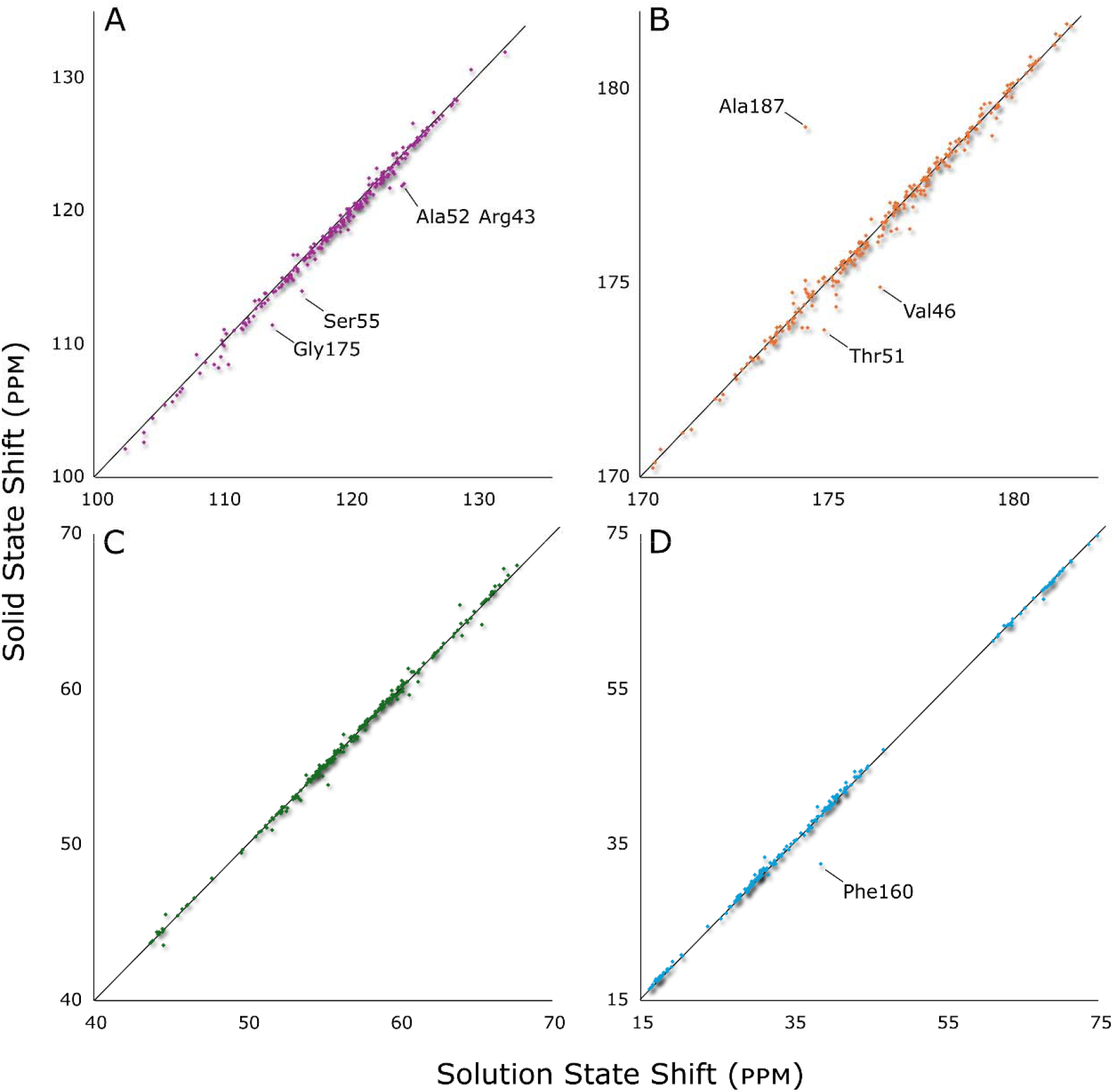
Comparison of backbone chemical shifts measured by solution-state NMR and solid-state NMR for Toho-1: (A) amide nitrogen; (B) carbonyl carbon; (C) alpha carbon; and (D) beta carbon. The close correspondence between datasets supports structural consistency of Toho-1 under crystallization and in solution.^12^ Several outliers are observed and correspond to residues located in or near regions of increased conformational dynamics identified in solution.

## 4. Conclusion

This study serves two key purposes. First, it demonstrates the power of ultrahigh-field NMR, combined with advances in probe technology and pulse sequence design, to achieve unprecedented resolution in solid-state spectra of microcrystalline proteins – even those as large as Toho-1. These technical improvements mark a significant step forward for the field. Second, this work lays the foundation for future sidechain analysis of the Toho-1 active site in the solid state. The close agreement between solution- and solid-state chemical shifts supports the integration of findings across both modalities and enables meaningful comparisons with prior studies of Class A β-lactamases.

## 5. Acknowledgements

## Author Contributions

CGW made and prepared samples. SW packed samples into NMR rotors. SW and CMR collected data. CGW, SW, CMR and OAW performed backbone and sidechain assignment and chemical shift analysis. CGW, LJM, SW and CMR wrote and prepared manuscript. All authors read and approved the final manuscript.

## Data availability

Assignments are submitted and available through the Biological Magnetic Resonance Data Bank (http://bmrb.io), Accession Number 53038.^38^

## Declarations

Authors declare no competing interests.

